# Potential virulence factors in *Pyrenophora teres* through label-free cellular proteomics analysis

**DOI:** 10.64898/2026.02.10.705184

**Authors:** Buddhika A. Dahanayaka, Sadegh Balotf, Richard Wilson, Anke Martin

## Abstract

*Pyrenophora teres* f. *teres (Ptt)* is the causative agent of net blotch diseases in barley and an economically important pathogen in the barley industry worldwide. To date, however, little is known about the protein expression profile of *Ptt*, which is important to understand the pathogen behaviour. In this study we report the first cellular proteomics analysis of *Ptt*. Label-free proteomics was used to quantify the protein expression levels of two parental and one of its progeny isolates from a *Ptt* cross, grown in culture. One parental isolate of the cross was virulent on the barley variety Prior while the other isolate was avirulent. The progeny isolate used in this study was also virulent on Prior. A total of 3,502 proteins were identified in samples of the three *Ptt* isolates, of which 99 were found only in the pathogenic isolates, while another 255 proteins were significantly more abundant in the pathogenic isolates compared to the non-pathogenic isolate. Gene ontology analyses of the significant proteins revealed that the proteins increased in pathogenic isolates were involved in fatty acid elongation, biosynthesis of unsaturated fatty acids, glycerophospholipid metabolism, nucleocytoplasmic transport, amino sugar and nucleotide sugar metabolism and metabolic pathways. These protein profiles and the bioinformatic analysis provide new biological information that can be utilised to better understand the pathogenicity of *Ptt*.

## Introduction

Net-form net blotch (NFNB) of barley (*Hordeum vulgare*) is an economically important disease wherever barley is grown. The yield losses due to NFNB were estimated to be up to 40% in susceptible cultivars (Murray & Brennan, 2010). In the absence of suitable fungicide treatment and during favourable environmental disease conditions, yield losses can increase to 70% (Wallwork et al., 2016). *Pyrenophora teres* f. *teres* (*Ptt*) [anamorph Drechslera (Sacc.) Shoemaker] is the causal agent of NFNB which is a foliar fungus belonging to the phylum Ascomycota. *Pyrenophora teres* f. *teres* is a highly variable pathogen that occur as a range of pathotypes from avirulent to virulent on different barley genotypes (Boungab et al., 2012; Oğuz & Karakaya, 2017; Steffenson & Webster, 1992). Previous studies have elucidated that different populations of *Ptt* collected globally are genetically diverse (Dahanayaka et al., 2021; Taliadoros et al., 2023) with multiple pathotypes having been observed.

Multiple QTL mapping studies have been conducted to understand the virulence mechanisms of *Ptt*. These studies have utilised multiple mapping methods such as bi-parental (Beattie et al., 2007; Dahanayaka et al., 2022; Koladia et al., 2017; Lai et al., 2007; Weiland et al., 1999), genome wide association (GWAS) (Martin et al., 2020) and multi-parental nested-association mapping (MP-NAM) (Dahanayaka & Martin 2023) populations to detect QTL. QTL were detected across the 12 *Ptt* chromosomes. The GWAS and MP-NAM studies have reported QTL mainly on chromosomes 3 and 5.

*Ptt* spends most of its lifecycle as a necrotrophic fungus acquiring nutrients from dead plant tissues (Able, 2003; Lightfoot & Able, 2010). The infection process of plant pathogens such as *Ptt*, is mainly facilitated by the physiological growth of the fungus and secretion of phytotoxins (Tan et al., 2010). During favourable temperature and humidity conditions the infection process of *Ptt* on susceptible barley cultivars initiates within six hours of the conidia/ascospores landing on the leaf lamella (Lightfoot & Able, 2010; Lightfoot et al., 2017). During the asymptomatic or short biotrophic phase of infection of *Ptt*, the germination tube of the conidia penetrates the leaf epidermal cells and then colonises the underlying mesophyll cells within 48h after inoculation. During its short biotrophic phase, *Ptt* produces pathogenesis-related (PR) proteins which are responsible for the activation of the salicylic acid (SA) pathway in the host (Al-Daoude et al., 2018). The activation of the SA pathway is a defence mechanism that the host adapts as a response to biotrophic fungi (Bauters et al., 2021; Bhadauria et al., 2010; Tsuda et al., 2008). The SA pathway causes cell death in the host by inducing the hypersensitive response (HR) to barricade the further spread of the pathogen (Bauters et al., 2021; Tsuda et al., 2008). However, cell death caused by activation of the SA pathway facilitates the growth of necrotrophic fungi (Faris et al., 2010; Friesen et al., 2007; Hammond-Kosack & Rudd, 2008) such as *Ptt* which exclusively relies on host cell death.

Previous studies using the secretome analysis of *Ptt* from culture filtrates identified a number of proteins that could induce necrotic symptoms on barley (Ismail & Able, 2016; Ismail et al., 2014a, 2014b; Sarpeleh, 2008; Sarpeleh et al., 2007). Proteins related to endoxylanase protein family (PttXyn11A), common in fungal extracellular membrane (CFEM), domain-containing protein (PttGPI) and cysteine hydrolase family protein (PttCHFP1) were identified in the culture filtrate of a virulent *Ptt* isolate, while they were absent in non-pathogenic *Ptt* isolates (Ismail et al., 2014a, 2014b). Proteomic analysis of culture filtrates from 28 *Ptt* isolates using two-dimensional gel electrophoresis (2-DE) identified 259 proteins with potential roles in the pathogenicity of *Ptt* (Ismail & Able, 2016). The majority of the proteins identified were associated with cell wall degrading enzymes, proteolytic, virulence effectors, proteins associated with fungal pathogenesis and development, and proteins related to oxidation–reduction processes.

Despite these insights from analysis of the secretome of different *Ptt* isolates, a comprehensive analysis of *Ptt* at the cellular proteome level has yet to be undertaken. Therefore, in the current study we used, a data independent acquisition mass spectrometry (DIA-MS) approach to compare a pathogenic *Ptt* isolate with its pathogenic and non-pathogenic parents to better understand the protein profile associated with *Ptt* virulence. This study is the first study to use large scale shotgun proteomics to report the proteomics profile of *P. teres*.

## Materials and Methods

### Biological materials

In this study we used two parental isolates (*P451* and *P155*) of a cross and a progeny isolates *SP23* (Dahanayaka, 2026). The crosses were developed as described in (Dahanayaka et al., 2022). Isolates *P451* and *P155* are progeny from a cross (P1) between *Ptt* isolates NB81 and HRS09127 used in the *Ptt* MP-NAM study (Dahanayaka & Martin 2023). Isolate *P451* is virulent (disease reaction score >6) on the barley cv. Prior and possesses a single QTL on chromosome 5 for Prior virulence while *P155* is avirulent (disease reaction score <6) on Prior and did not possess any QTL associated with Prior virulence (Figure 1).

**Figure 1.**
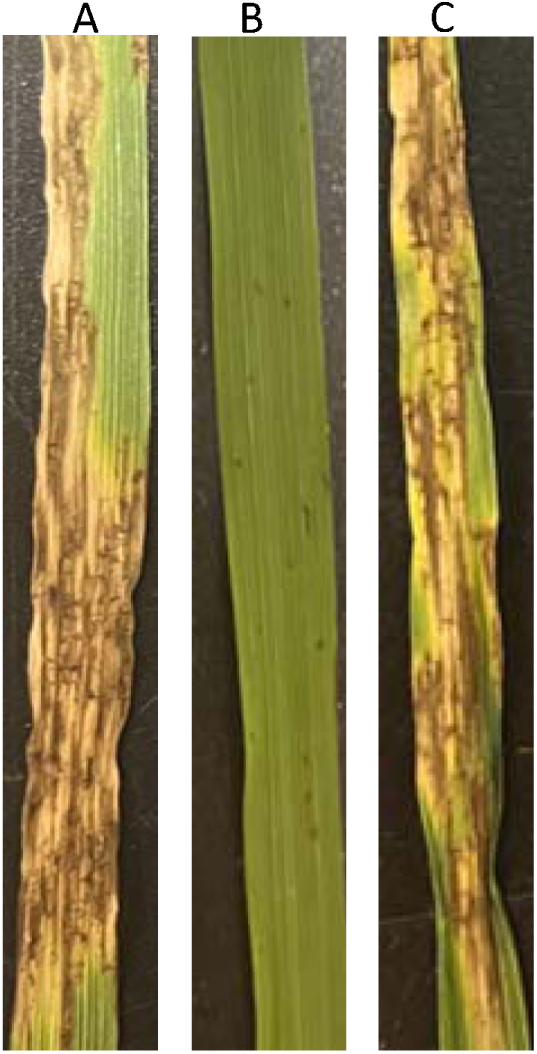
Disease symptoms caused by three *P. teres* f. *teres* isolates A. virulent (*P451*) B. avirulent (*P115*) and C. progeny (*SP23*) on barley cv. Prior.

### Protein extraction

Isolate *SP23* is a progeny of the cross between *P451* and *P155* and is pathogenic on Prior. The three *Ptt* isolates, *P451, P155* and *SP23* were grown on PDA medium at 22°C for 10 days. Mycelium from each isolate was scraped from the PDA medium and freeze-dried for 24 hours. The freeze-dried mycelium from the *Ptt* isolates was used for protein extraction.

Protein extraction was conducted as three independent replicates for each isolate grown on PDA. Proteins were extracted from 50-70 mg of frozen samples by adding 150 µL of extraction buffer [7M urea, 2M thiourea, 1% dithiothreitol (DTT), 100mM NaCl, 40 mM Tris, pH8.0]. The samples were homogenised for 30 s using a Fast Prep-24 bead beater (Mp Biomedicals, Seven Hills, NSW, Australia) at 5500 rpm at room temperature and then centrifuged at 16,000 rpm for 10 min at 4°C. The supernatant was transferred to a 1.5mL tube. Six volumes of cold acetone were added to the resulting clarified solution and samples were incubated overnight at −20°C. The samples were centrifuged at 10,000 rpm for 8 min at 4 °C. The resultant protein pellets were left to air dry for 5 min at room temperature before resuspending in the same extraction buffer mentioned above.

### Protein reduction, alkylation and digestion

Samples were reduced overnight at 4°C using DDT at the final concentration of 10 mM. Subsequently, the reduced proteins were alkylated by adding iodoacetamide to each sample at the final concentration of 50 mM, followed by 2 h incubation at room temperature in the dark. The Single-Pot Solid-Phase-enhanced Sample Preparation (SP3) method (Hughes et al., 2019) was used for the trypsin/LysC digestion of the proteins. The resulting peptides were then desalted using ZipTips (Merck, Darmstadt, Germany).

### Data acquisition, raw data processing and bioinformatic analysis

Desalted peptides were dried in a SpeedVac concentrator and dissolved in 12 µL 0.05% (w/w) trifluoroacetic acid in water/acetonitrile (98:2). One μg of peptide per sample was analysed using an Ultimate 3000 nano RSLC system coupled to an Orbitrap Q-Exactive HF mass spectrometer. Samples were separated over 2 h LC gradients and analysed using data-independent mass spectrometry (DIA-MS) according to the parameters described previously (Balotf et al., 2021). Raw DIA-MS files were imported into Spectronaut software (v 17, Biognosys) and searched against the UniProt *Pyrenophora teres* protein sequence database (UP000001067) accessed on March 3^rd^ 2023 and comprising 11,705 entries. Spectronaut software was then used for targeted DIA analysis of the resulting spectral library. Default Biognosys (BGS) settings were used for both library generation and extraction of MS1 and MS2 DIA profiles, with the exception that single-hit protein ID’s were excluded from the quantitative analysis.

The protein intensity values were filtered by removing proteins with more than one missing value per isolate. The protein intensity values were then transformed (log_2_) and any remaining missing values were imputed from the normal distribution using Perseus v2.0.9.0 (Tyanova et al., 2016) according to default settings. The principal component analysis (PCA) plots and heatmaps were obtained from MetaboAnalyst5 (Pang et al., 2021).

Proteins were classified as significantly altered in abundance between groups on the basis of the Student’s t-test according to a False Discovery Rate (FDR□<□0.05, with the s0 parameter set to 0.1. The association networks for the highly significant proteins were constructed using ShinyGO (Ge et al., 2020) and functional enrichment of the differentially abundant proteins was obtained from the STRING database (https://string-db.org/) based on a threshold of FDR <0.05.

Proteins that were detected only in the pathogenic parental and pathogenic progeny and not in the non-pathogenic parental isolate were identified manually in order to recognise potential proteins that could be important for the virulence of *P. teres*. These proteins and the proteins showed significant increase abundance in pathogenic parental and pathogenic progeny compared to the non-pathogenic parental isolate were used in EffectorP 3.0 (Sperschneider & Dodds, 2022) to detect potential effectors which could be important for the infection process of *P. teres*. Furthermore, proteins that were unique or showed more than 5-fold increase in abundance (*P*□≤□0.05) in pathogenic parental and pathogenic progeny isolates compared to the non-pathogenic isolate were located in the *P. teres* f. *teres* reference genome; W1-1 (BioSample:SAMEA4560035 and BioProject:PRJEB18107). Maps of these protein positions together with the *P. teres* f. *teres* virulence QTL reported to-date were produced using MapChart 2.23 (Voorrips, 2002).

## Results

### Overview of the protein profile of Ptt isolates

The proteome profiles of the pathogenic (*P451* and *SP23*) and non-pathogenic (*P155*) *Ptt* isolates were created using a DIA-MS approach in which a library of 13,995 peptide sequences mapping to a total of 3,800 protein groups was generated. After filtering single-hit proteins as well as proteins with more than one missing value per isolate, 2,343 proteins were retained for further analyses (Supplementary Table 1). Several hundred proteins were significantly altered in abundance between the two groups based on Student’s t-test (*P/FDR*□<□0.05) analysis. The complete list of proteins retained for further analyses with their relative abundance values is provided in Supplementary Table S1.

Principal component analysis (PCA) was used to display the relationships between non-pathogenic and pathogenic parental and progeny isolates of *Ptt*, indicating a clear separation among different *Ptt* isolates which accounted for more than 50% of the total variation (Figure 2). Hierarchical clustering of the total protein set and representation of Z-scored protein intensity values as a heat map also showed distinct clusters for each isolate (Figure 3). Consistent with the PCA plot, each of the samples were present in separate sub-clusters.

**Figure 2.**
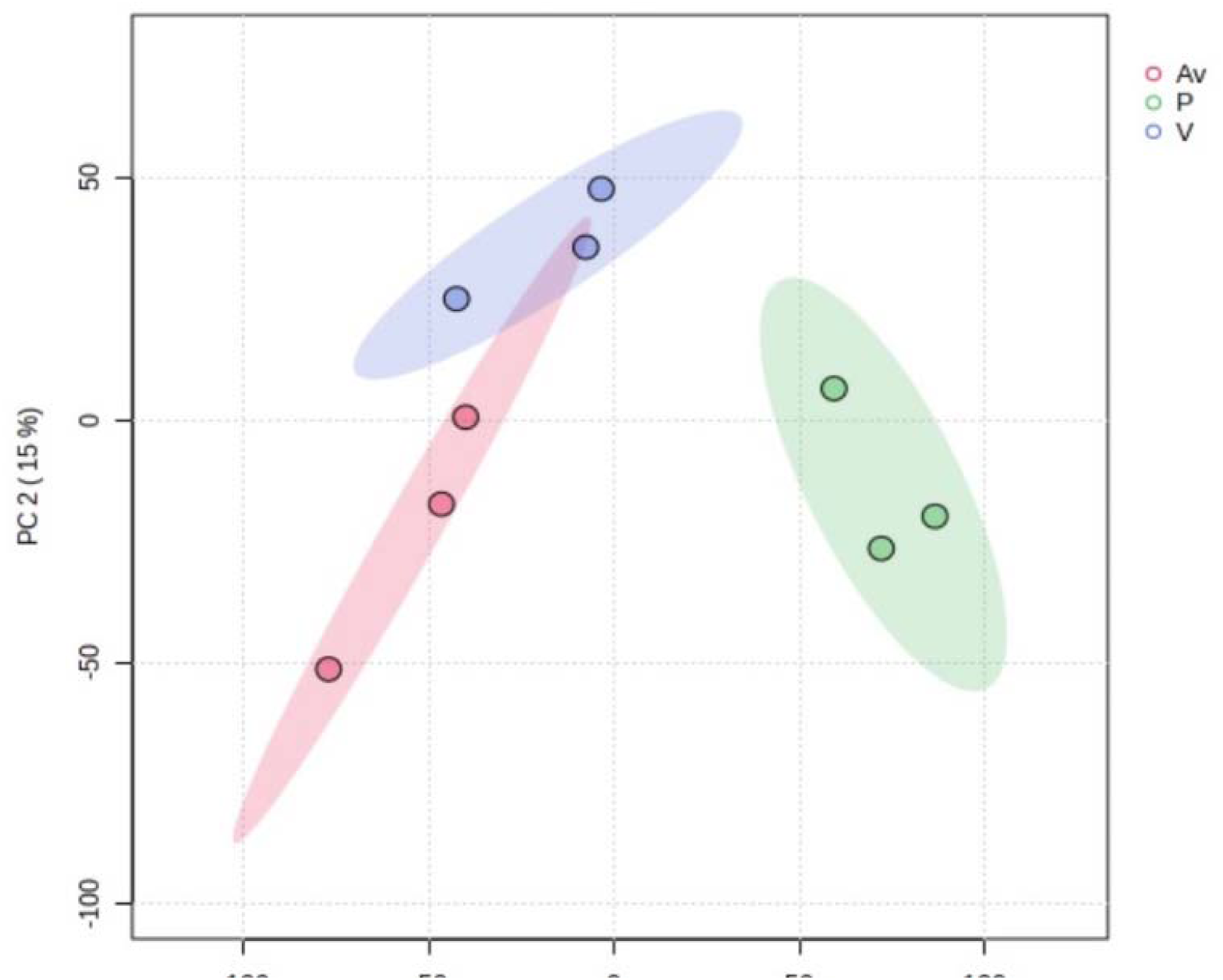
Principal Component Analysis (PCA) showing the variations in the whole proteomes of pathogenic and non-pathogenic parental *Pyrenophora teres* f. *teres* isolates, as well as a pathogenic progeny isolate. Proteomics analysis was performed with three biological replicates per isolate. Av: non-pathogenic parent (*P155*); P: pathogenic progeny (*SP23*); V: pathogenic parent (*P451*)

**Figure 3.**
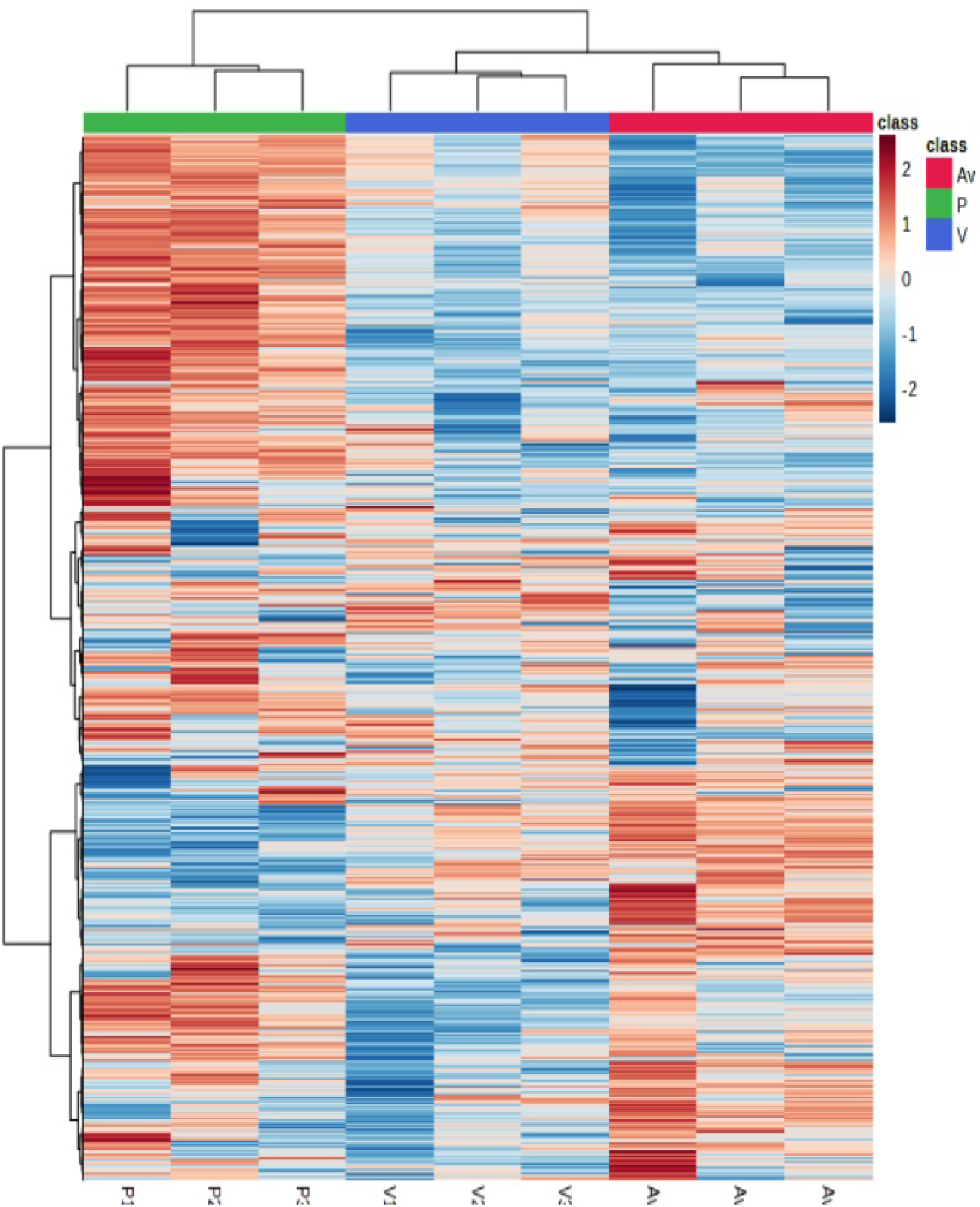
Heatmap plot of standardized abundance values for the full set of proteins represented as Z-scored intensity values. Av: non-pathogenic parent (*P155*); P: progeny (*SP23*); V: pathogenic parent (*P451*)

Comparisons among protein profiles of the isolates were used to identify significantly abundant proteins for each isolate. Comparison of the pathogenic progeny (*SP23*) isolate with the pathogenic (*P451*) and non-pathogenic (*P155*) parental isolates using t-test analysis (Figure 4A and B, respectively) identified 753 and 886 proteins, respectively, that were significantly changed in abundance (Supplementary Tables S1 & S2). Out of these sets of significant proteins, 236 and 351 proteins were altered by ≥3-fold in *SP23* samples compared to the *P451* and *P155* samples, respectively (Supplementary Tables S2 & S3). Consistent with the PCA and cluster analysis, fewer significant proteins were identified by t-test comparison of the pathogenic (*P415*) and non-pathogenic (*P155*) parental isolates (Figure 4C), in which 33 proteins were significantly changed, of which 11 proteins were altered by ≥3-fold (Supplementary Table S4).

**Figure 4.**
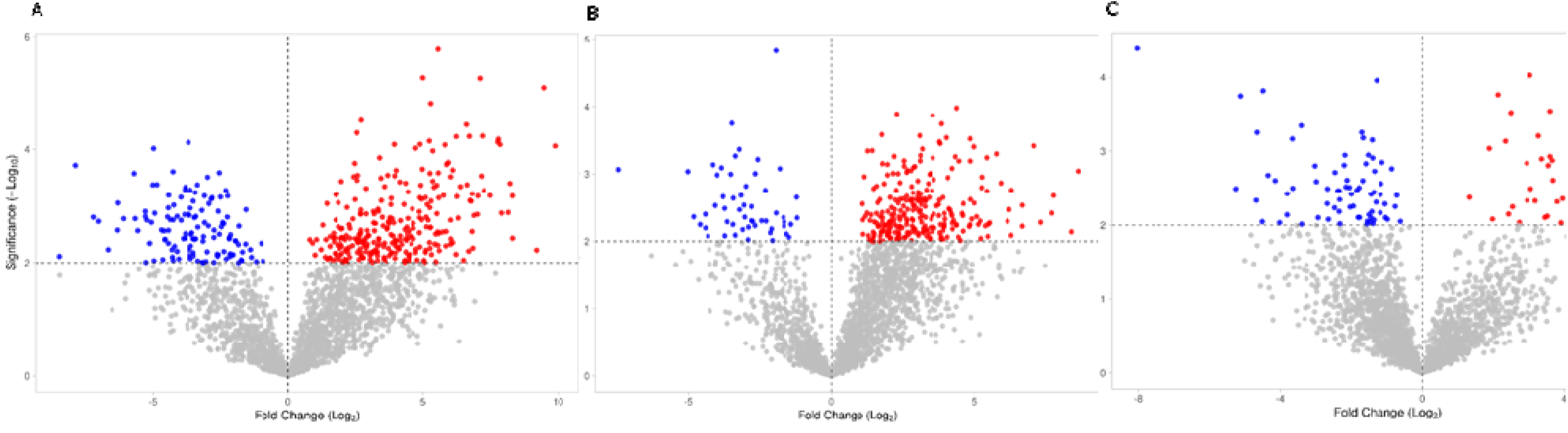
Volcanic plots showing significantly increased (red) and decreased (blue) proteins (FDR□≤□0.05) of the *Pyrenophora teres* f. *teres* progeny isolate (*SP23*) compared to the non-pathogenic (*P155*) (A) and pathogenic (*P451*) parent (B) and the pathogenic parent compared to the non-pathogenic parent (C). The x-axis and the y-axis represent the magnitude of the fold-changes and the statistical significance (-log10 of p-values) of the protein values. The dashed horizontal and vertical lines denote the cut-off for p-values and proteins that are either significantly increased or decreased in abundance. Grey points represent proteins which did not show a significant increase or decrease in abundance.

To identify the proteins which may contribute for the virulence of *P. teres*, the protein profiles of the pathogenic isolates and the non-pathogenic isolate were compared. The protein profiles of the pathogenic isolates *P451* and S*P23* reported 511 differentially abundant proteins (*P* <□0.05) compared to the non-pathogenic isolate *P155*. Out of these 511 proteins, 255 showed significant increased abundance in the pathogenic isolates compared to the non-pathogenic isolate (Table 1 and Supplementary Table S5) and some of these had previously been characterised in other micro-organisms (Table 1). Another 99 proteins out of 2,343 were found to be present only in the pathogenic isolates and absent in the non-pathogenic parental isolate (Supplementary Table S6).

**Table 1.** List of proteins showing greater than 5-fold increase in abundance (*P*□≤□0.05) in the pathogenic isolates compared to the non-pathogenic isolate of *Ptt*. Proteins that have previously been identified in other studies and organisms are reported.

| Gene ID | Protein ID | Protein description | Increased fold change | Organisms | Reference |
| --- | --- | --- | --- | --- | --- |
| *PTT_16464* | E3S2D9 | Steroid 5-alpha reductase C-terminal domain-containing protein | 8 | NA | NA |
| *PTT_10048* | E3RNA8 | Golgi apparatus membrane protein TVP23 | 8 | *Magnaporthe oryzae* | Zhao et al. (2022) |
| *PTT_12687* | E3RUD5 | Derlin | 6 | NA | NA |
| *PTT_16451* | E3S2C8 | Cleft lip and palate associated transmembrane protein 1 | 6 | NA | NA |
| *PTT_16825* | E3S337 | Deacetylase sirtuin-type domain-containing protein | 6 | *Ustilago maydis*<br>*Fusarium oxysporum* | Navarrete et al. (2023); Zhao and Rusche (2022); Zhang et al. (2023) |
| *PTT_03618* | E3RE30 | Uncharacterized protein | 5 | NA | NA |
| *PTT_08158* | E3RJ63 | Anoctamin/TMEM 16 | 5 | NA | NA |
| *PTT_09405* | E3RLY2 | KRR1 small subunit processome component | 5 | *Fusarium*<br>*oxysporum*<br>*Valsa mali* | Nandhini et al.<br>(2018)<br>Xu et al. (2020) |
| *PTT_10832* | E3RQ68 | Zf-C3HC-domain-containing protein | 5 | NA | NA |
| *PTT_18618* | E3S748 | Transmembrane protein UsgS | 5 | NA | NA |
| *PTT_02290* | E3RDH9 | Chitin synthase | 5 | *Fusarium*<br>*oxysporum*<br>*Magnaporthe*<br>*oryzae*<br>*Cryptococcus*<br>*neoformans*<br>*Phytophthora*<br>*capsici and*<br>*Phytophthora sojae* | Madrid et al. (2003)<br>Kong et al. (2012)<br>Banks et al. (2005)<br>Cheng et al. (2019) |
| PTT_12879 | E3RUT9 | Polynucleotide adenylyltransferase | 5 | NA | NA |
| PTT_15408 | E3S059 | ER membrane protein complex subunit 4 | 5 | NA | NA |
| PTT_17997 | E3S5R0 | Mechanosensitive ion channel protein | 5 | NA | NA |
| PTT_18673 | E3S796 | CaaX prenyl proteinase Rce1 | 5 | NA | NA |
| PTT_16864 | E3S364 | Protein transport protein SEC22 | 5 | *Magnaporthe*<br>*oryzae*<br>*Colletotrichum*<br>*orbiculare* | Song et al. (2010)<br>Irieda et al. (2014) |

The 99 proteins, with the exception of two, that were only present in the pathogenic isolates were located on the *P. teres* f. *teres* reference genome, W1-1 together with previously reported *Ptt* virulence QTL, showing that some of these are co-located (Figure 5). Fourteen proteins that had a greater than 5-fold increase in abundance (P□≤□0.05) in the pathogenic parental and progeny isolates compared to non-pathogenic isolates were also co-located with previously identified QTL (Figure 5).

**Figure 5.**
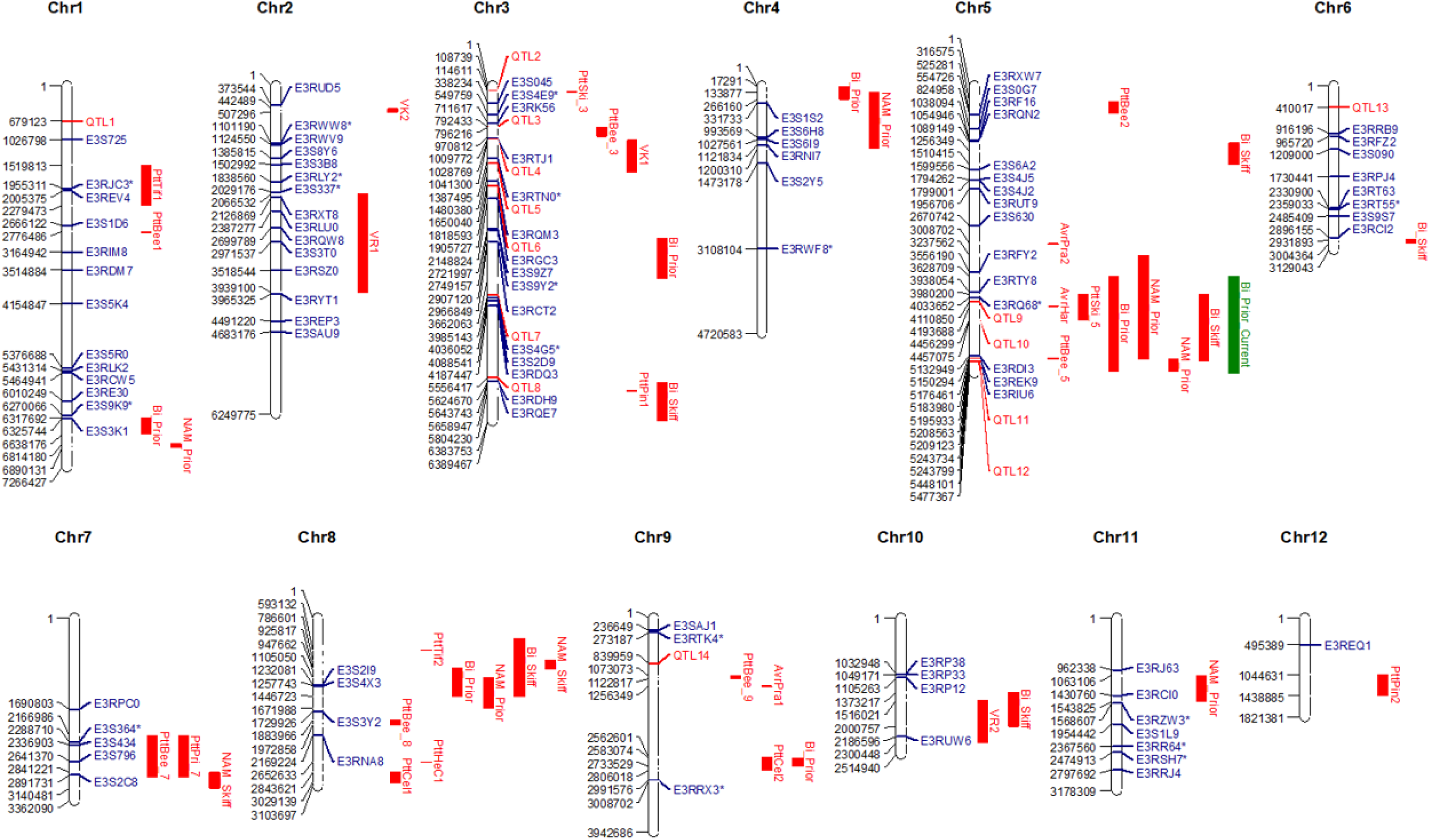
Map of *P. teres* f. *teres* reference genome, W1-1. Location of proteins only expressed or showed 5-fold increase abundance (P□≤□0.05) in the pathogenic isolates compared to the non-pathogenic isolate are indicated in blue. Previously reported QTL and QTl identified from the current study associated with *P. teres* f. *teres* virulence are indicated in red (Dahanayaka et at. 2023 and Martin et al, 2021) and green, respectively. Effector proteins as predicted by EffectorP 3.0 are designated with an asterisk.

### Annotation and gene ontology enrichment analysis

Using gene ontology (GO) analysis, sets of significantly abundant proteins were categorised into groups based on their molecular function (MF), cellular component (CC), and biological process (BP). The sets of significantly abundant proteins of the pathogenic progeny isolate *SP23*, compared to the pathogenic and non-pathogenic parents identified 20 and 16 significantly enriched proteins (FDR<0.05) were used for functional enrichment analysis (Figures 6 and 7). Most of these proteins were involved in the biosynthesis of secondary metabolites and metabolic pathways.

**Figure 6.**
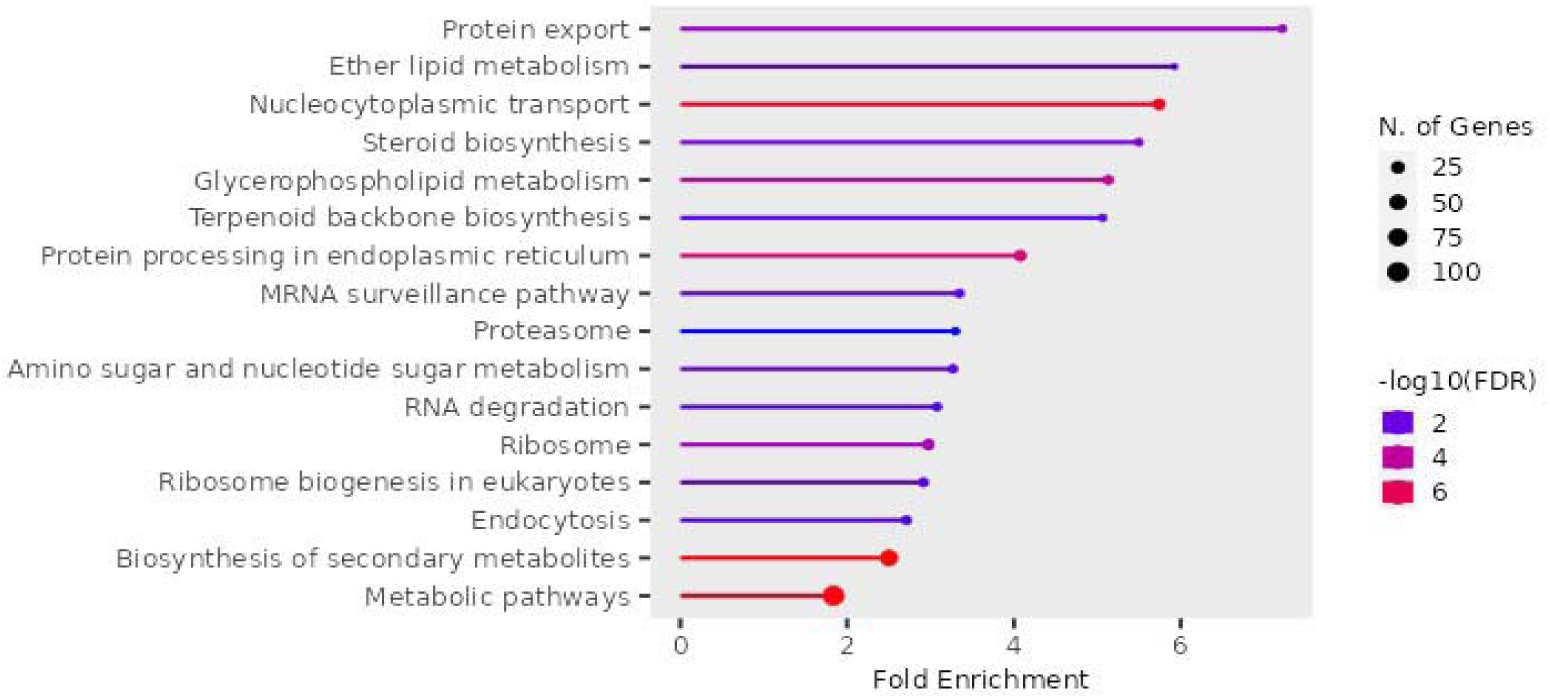
Functional enrichment analysis of the increased differential protein abundance in the pathogenic progeny isolate compared to the non-pathogenic parental isolate of *Ptt*.

**Figure 7.**
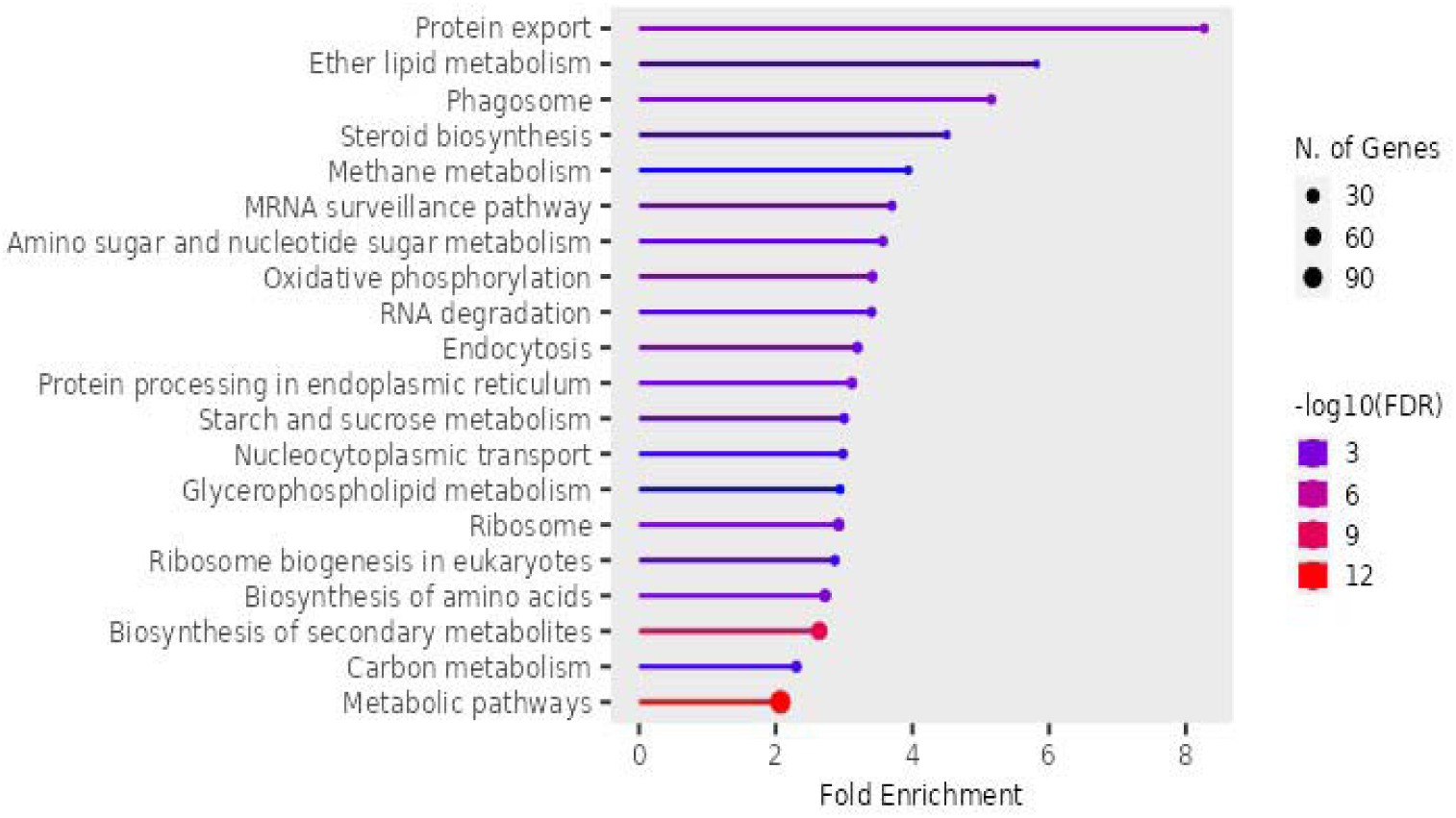
Functional enrichment analysis of the increased differential protein abundance in the pathogenic progeny isolate compared to the pathogenic parental isolate of *Ptt*.

Proteins that were only expressed in the pathogenic isolates (99 proteins) belonged to six functional protein categories, namely fatty acid elongation, biosynthesis of unsaturated fatty acids, glycerophospholipid metabolism, nucleocytoplasmic transport, amino sugar and nucleotide sugar metabolism and metabolic pathways (Figure 8).

**Figure 8.**
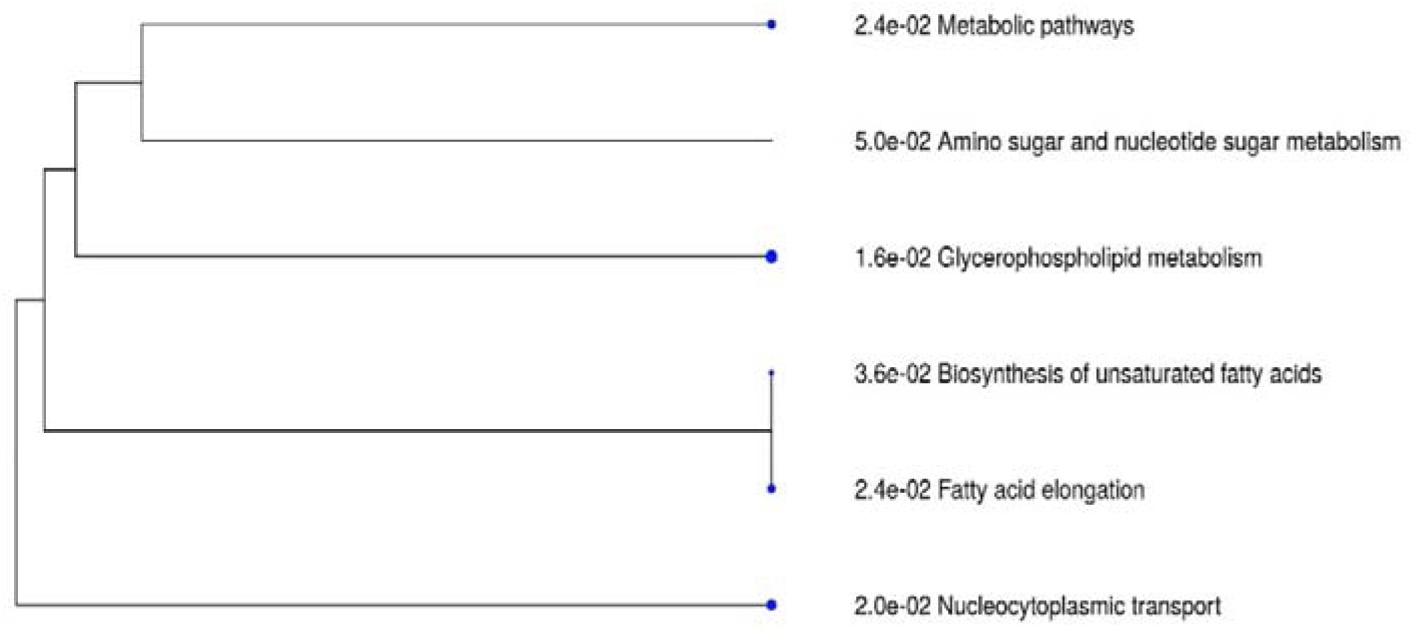
A hierarchical clustering tree illustrating the correlation among the pathways showing increased differential protein abundance in pathogenic isolates compared to non-pathogenic *Ptt* (with a p-value < 0.05 and a log2 fold change > 2). Pathways sharing multiple genes are grouped together, and larger blue dots denote more significant p-values.

Effector mining of the proteins that were only expressed and showed significant increase in abundance in the pathogenic isolates compared to the non-pathogenic isolates detected 175 candidate effectors that could be important for the virulence of *P. teres* (Figure 5 and Supplementary Table S7). The majority (155) of the predicted effectors were cytoplasmic effectors and 13 were apoplastic effectors. Another four of the effectors were cytoplasmic/apoplastic effectors and the other three were apoplastic/cytoplasmic effectors.

## Discussion

This study reports the first LC-MS protein profiling of *P. teres*. The protein profile of a pathogenic and non-pathogenic *Ptt* isolates were compared using large-scale shotgun proteomics. Hundreds of significantly increased abundant proteins were detected and a comprehensive protein profile established. The co-locations of these proteins with previously identified *P. teres* f. *teres* virulence QTL using the W1-1 reference genome is a valuable resource for other researchers to quickly identify proteins potentially important in the virulence of *P. teres*.

In *in planta* proteomics studies of fungal pathogens, the detection of fungal proteins is limited due to the predominance of host plant proteins and the relatively low biomass of the fungal pathogen (González-Fernández et al., 2010). For example, in a study on *Fusarium graminearum*, high-throughput MS/MS was used to identify proteins secreted during growth on various media *in vitro* and *in planta* during wheat head infection. A total of 289 proteins were identified, with 229 *in vitro* and 120 *in planta* (Paper et al., 2007). A study on biotrophic barley fungal pathogen *Blumeria graminis* f*. sp. hordei* found differences between *in vitro* and *in vivo* protein conditions. The study identified 441 proteins in ungerminated spores, 775 proteins in epiphytic sporulating hyphae, and 47 proteins from haustoria inside barley leaf epidermal cells (Bindschedler et al., 2009). Due to the limited detection of fungal proteins in *in planta* studies, the current research was conducted on fungal cultures grown in media to identify as many fungal proteins as possible from the cellular level of the fungal isolates.

A significant increase in abundance of 255 proteins was observed with the pathogenic isolate protein profiles compared to the non-pathogenic isolate. Another 99 proteins only expressed in the pathogenic isolates may play an important role in *P. teres* virulence. Some of these proteins, either showing a significant increase in abundance or expressed only in pathogenic isolates, were co-located with previously identified QTL.

Previously reported QTL on chromosome 5 associated with virulence on genotype Prior and the QTL identified in the current study (Dahanayaka & Martin 2023; Martin et al., 2020) co-located with six proteins (E3RFY2, E3RTY8, E3RQ68, E3RDI3, E3REK9 and E3RIU6). Two of these were uncharacterised proteins (E3RFY2 and E3RIU6) and the other four proteins were E3RTY8; Heterokaryon incompatibility (HET) domain-containing, E3RQ68; ZfC3 HC-domain-containing protein, E3RDI3; Benzoate 4-monooxygenase cytochrome P450 and E3REK9; Chitin synthase. Studies have shown that HET domain-containing proteins are involved in the self-recognition system of filamentous fungi. These proteins help prevent genetically distinct fungal strains from forming viable cell fusions, or heterokaryons, by triggering programmed cell death when their HET loci are incompatible (Paoletti & Clave, 2007; Saupe, 2000; Saupe et al., 2000). Liu et al. (2020) have shown that HC-domain-containing proteins are play a major role in the regulation of genes, stress responses, integrity of the cell wall, conidiation, development of infection hyphae and MAP1-mediated pathogenicity in *Magnaporthe oryzae*. Zhang et al. (2012) detected increased expression of a cytochrome enzyme Benzoate 4-monooxygenase in *Fusarium graminearum* during its invasion of wheat coleoptiles, suggesting that this protein plays a role in pathogenicity. A previous study has shown that when the genes that are responsible for chitin synthase are impaired, the conidial germination and production are negatively affected and the virulence in *Verticillium dahlia* is reduced (Qin et al., 2022).

Our proteomic analysis of *Ptt* isolates also detected proteins in QTL regions that were not targeted in this fungal population. Presence of proteins/candidate effectors in non-targeted regions suggest that there may be additional virulence factors or other functional proteins within the genome of these isolates, which could play a role in the pathogen’s interaction with hosts other than Prior. Previously reported QTL mapping studies using bi-parental (Martin et al., 2019) and nested association (Dahanayaka & Martin 2023) populations of *P. teres* f. *teres* detected significant QTL responsible for Prior, Beecher and Skiff virulence on chromosome 7 and three proteins (E3S364, E3S434 and E3S2C8) of the current study, which were only expressed in pathogenic isolates, were also found in the same region. E3S364; transport protein SEC22, which was predicted to be an effector by the EffectorP, was found to be important for the integrity of the cell wall, pathogenicity, reproduction, regulation of reactive oxygen species and the expression of extracellular enzymes in filamentous fungi (Adnan et al., 2019). E3S434; Phox homology (Px) domain containing protein was shown to be important in plant infection and deoxynivalenol (DON) production in *Fusarium graminearum* (Lou et al., 2021). E3S2C8; cleft lip and palate associated transmembrane protein 1 is responsible for the cleft lip birth defect in humans. This protein was also found to be present in plants and other fungi, but the function of the protein has not been determined as per authors knowledge. Future studies to demonstrate the function of these proteins in *barley-P. teres* pathosystem will be important.

A significantly increased abundance of various ATP-binding cassette (ABC) transporter proteins was detected in the pathogenic isolates compared to the non-pathogenic isolate in the current study. It has previously been reported that ABC transporter proteins regulate pathogenicity in plant fungal pathogens (Idnurm & Howlett, 2001). Mutations of proteins in ABC transporter families, ABC1 (Urban et al., 1999) and ABC4 (Gupta & Chattoo, 2008) of *Magnaporthe grisea,* resulted in the defective appressorium formation and thus the loss of pathogenicity. Disruption of *FgABCC9*, an ABC transporter protein in *F. graminearum* exhibited low levels of disease reactions in wheat (Qi et al., 2018). ABC transporter proteins also play a major role in fungicide resistance in filamentous fungi. Knockout mutants of *MoABC-R1* gene in *M. oryzae* showed significantly increased sensitivity to pyraclostrobin (Hu et al., 2023).

The pathogenic isolates of the current study compared to the non-pathogenic isolate demonstrated a significantly increased abundance of serine/threonine protein kinases (PK). Serine/threonine PK catalyses the phosphorylation of serine/threonine residues on a target protein by using ATP as phosphate donor and may results in changes in the function of the target protein. Mitogen-activated protein kinase (MAPK) is a group of serine/threonine kinase. *PTK1* a MAPK gene in *P. teres* was found to be required for appressorium formation and conidiation (Ruiz-Roldán et al., 2001) with *P. teres* mutants of *PTK1* producing abnormally elongated conidiophores without conidia formation. Similar observation was also reported in *CHK1* mutants with *Cochliobolus heterostrophus* where the mutants were unable to form conidia (Lev et al., 1999) or had reduced conidia production such as *CMK1* mutants of *C. lagenarium* (Takano et al., 2000) and *MPS1* mutants of *M. grisea* (Xu et al., 1998). MAPK (*CsSTE11,* and *CsFUS3*) mutants of *Bipolaris sorokiniana,* another cereal pathogen, also showed a significant reduction in virulence compared to its wild type (Leng & Zhong, 2015). Serine/threonine protein kinase in *M. oryzae* (*MoSch9*) was also reported to regulate the sporulation and pathogenesis of the fungus (Chen et al., 2014). The mutant *Mosch9* resulted in smaller conidia and appressoria, reducing the hyphal growth and pathogenesis (Chen et al., 2014).

A previous study based on proteomics analyses of *P. teres* detected 259 individual proteins from culture filtrates of 28 *Ptt* pathogenic isolates (Ismail & Able, 2016). A subsequent study aimed at detecting the expression of 222 genes out of 259 previously identified proteins, discovered high to extremely high expression levels in virulence proteins contributing to oxidation-reduction, or carbohydrate metabolic processes (CMP) including the cell wall degrading enzymes (CWDEs) (Ismail & Able, 2016). In our study, 119 proteins of those reported in the previous *P. teres* proteomics study (Ismail & Able, 2016) were detected, 49 out of which were increased in their abundance (Supplementary Table S5) in the pathogenic isolates compared to the non-pathogenic isolate. Both studies consistently found high expression levels of key virulence proteins, highlighting their crucial role in the organism’s pathogenicity and potential as targets for the disease.

The PCA plot and the heatmaps of the expression data indicated that the protein profiles of the non-pathogenic and pathogenic parental isolates were more closely related to each other than to the pathogenic progeny isolate. The parental isolates are genetically closely related, except for the genomic regions associated with virulence, as they have been derived from the same cross. Even though the genes which are responsible for the pathogenicity of the progeny isolate would have been acquired from the pathogenic parent of the cross, the protein profile of the progeny isolate showed a different abundance in its protein profile compared to that of the pathogenic parent. While proteomic analysis of fungal cultures revealed differences in protein expression between the pathogenic progeny and the virulent parent isolate, phenotyping indicated that they exhibit similar pathogenic behaviour. This suggests that the presence of the host may influence pathogen gene expression and protein activity in ways not captured *in vitro*. Therefore, to better understand the molecular basis of pathogenicity, future work should include in planta experiments to capture the host-induced dynamics of protein expression.

This first protein profile of *P. teres* using LC-MC identified 255 proteins that were significantly increased in their abundance in the pathogenic isolates compared to the non-pathogenic isolate and another 99 proteins that were present only in the pathogenic parental and progeny isolate and absent in the non-pathogenic parental isolate. Some of these proteins had been characterised by previous studies as potential virulence factors. This study has provided insights into the proteins that *Ptt* may potentially produce during infection. Future studies to identify and validate the function of more of these proteins specifically proteins that were recorded in previous QTL mapping studies would be useful for candidate gene identification.

## Data Availability Statement

The mass spectrometry proteomics data have been deposited to the ProteomeXchange Consortium via the PRIDE (Perez-Riverol et al., 2019) partner repository with the dataset identifier PXD050290. Project Name: Label-free proteomics analysis of *Pyrenophora teres* isolates reveals potential virulence factors. Username. Password: iYVPjX7X

## Author contributions

B.D. and S.B. performed the experiments. R.W. performed the initial proteomics analyses. B.D. performed the analyses and prepared and revised the manuscript. S.B., R.W. and A.M. reviewed the manuscript.

## Ethics declarations

The authors declare no competing interests.

